# Use of an individual-based model of pneumococcal carriage for planning a randomized trial of a vaccine

**DOI:** 10.1101/258871

**Authors:** Francisco Y. Cai, Thomas Fussell, Sarah E. Cobey, Marc Lipsitch

## Abstract

For encapsulated bacteria such as *Streptococcus pneumoniae*, asymptomatic carriage is more common and longer in duration than disease, and hence is often a more convenient endpoint for clinical trials of vaccines against these bacteria. However, using a carriage endpoint entails specific challenges. Carriage is almost always measured as prevalence, whereas the vaccine may act by reducing incidence or duration. Thus, to determine sample size requirements, its impact on prevalence must first be estimated. The relationship between incidence and prevalence (or duration and prevalence) is convex, saturating at 100% prevalence. For this reason, the proportional effect of a vaccine on prevalence is typically less than its proportional effect on incidence or duration. This relationship is further complicated in the presence of multiple pathogen strains. In addition, host immunity to carriage accumulates rapidly with frequent exposures in early years of life, creating potentially complex interactions with the vaccine’s effect. We conducted a simulation study to predict the impact of an inactivated whole cell pneumococcal vaccine—believed to reduce carriage duration—on carriage prevalence in different age groups and trial settings. We used an individual-based model of pneumococcal carriage that incorporates relevant immunological processes, both vaccine-induced and naturally acquired. Our simulations showed that for a wide range of vaccine efficacies, sampling time and age at vaccination are important determinants of sample size. There is a window of favorable sampling times during which the required sample size is relatively low, and this window is prolonged with a younger age at vaccination, and in a trial setting with lower transmission intensity. These results illustrate the ability of simulation studies to inform the planning of vaccine trials with carriage endpoints, and the methods we present here can be applied to trials evaluating other pneumococcal vaccine candidates or comparing alternative dosing schedules for the existing conjugate vaccines.

**Author Summary:** *Streptococcus pneumoniae*, a bacterium carried in the nasopharynx of many healthy people, is also a leading cause of bacterial pneumonia, sepsis, and ear infections in children aged five years and younger. Vaccines targeting select strains of *S. pneumoniae* have been effective, and the development of new vaccines, particularly those that target all strains, can further lower disease burden. For clinical trials of these vaccines, the number of study participants needed depends on the expected effect of the vaccine on a conveniently measured outcome: asymptomatic carriage. The most economical way to test a vaccine for its effect on carriage is by measuring prevalence at a specific time, and comparing vaccinated to unvaccinated participants. The relationship between incidence (or duration) and prevalence is complex, and changes with time as children develop natural immunity. We explored this relationship using a mathematical model. Given a vaccine efficacy, our computer simulations predict that fewer study participants are needed if they are vaccinated at a younger age, taken from a population with intermediate levels of transmission, and sampled for carriage at a certain time window: 9 to 18 months after vaccination. Our study illustrates how simulation studies can help plan more efficient vaccine trials.

## Introduction

For encapsulated bacteria such as *Streptococcus pneumoniae* [1], *Haemophilus influenzae* [2], and *Neisseria meningitidis* [3], asymptomatic carriage in the human upper respiratory tract is a precursor to mucosal or invasive disease. The population of bacteria in the upper respiratory tract, which may be sampled in the oropharynx or nasopharynx, is also the primary or sole source of transmission of these bacteria. Because carriage is far more common and typically longer in duration than disease with these bacteria, it is often a more convenient endpoint for clinical trials of vaccines against them. If a vaccine can prevent or terminate carriage, then it is likely to reduce both the risk of disease and the opportunities for transmission, leading to herd immunity effects. Many of the current generation of vaccines against these organisms, made from their capsular polysaccharides chemically conjugated to a protein carrier (conjugate vaccines), have been evaluated in randomized controlled trials (RCTs) where carriage was the primary endpoint [4–10], and the case for carriage as an endpoint in vaccine licensure has been put forth by an international consortium [11]. Carriage endpoints have also been used for RCTs of other vaccines against encapsulated bacteria, such as the protein-based vaccine designed to protect against group B meningococci [12].

While the use of carriage as an endpoint in an RCT is often convenient and offers the possibility of smaller sample sizes than disease endpoints, it presents added complexities. Carriage is almost always measured as prevalence (whether the target organism is present at a particular time) rather than as incidence (the rate at which individuals acquire the organism), the more traditional endpoint in vaccine trials. For vaccines such as conjugate vaccines that are thought to act directly on vaccinated persons by reducing the incidence of acquiring colonization, the proportional reduction in prevalence due to a vaccine will in general be smaller than the proportional reduction in incidence it causes [13], because prevalence increases less than linearly with incidence. Under certain assumptions, the estimated impact on prevalence can be converted into an estimate of the impact on incidence [13], though this becomes more complex when there are multiple serotypes targeted by the vaccine [14]. At a practical level, decisions must be made about when to sample the carriage population to estimate efficacy, with the goal of observing the largest effect possible (to reduce sample size) and also of being able to estimate a meaningful efficacy parameter [15]. Moreover, immunity to carriage of S. *pneumoniae* (also called pneumococci, the species on which this paper and the remainder of this introduction will focus) likely involves at least two different parts of the immune system: antibodies that act in a serotype-specific fashion to reduce the risk of acquisition [16] and T-helper cells that act in a serotype-independent manner to reduce the duration of a carriage episode [17]. Both of these forms of immunity are imperfect: even after multiple exposures to pneumococci, a human can acquire colonization and will not clear it immediately [16,18,19]. Vaccines typically augment or hasten the acquisition of immunity, but vaccine-induced immunity against carriage is also only partially effective [13]. In a vaccine trial conducted in infants or toddlers, participants in both the vaccine group and the control group will be repeatedly challenged by exposure to pneumococci. Through the experience of acquiring and clearing colonization, these individuals will develop immune responses that reduce their rate of acquisition on exposure and increase the rate at which they clear the colonization episode [16,20]. Further complexity arises from the fact that individuals may be colonized simultaneously with multiple strains of pneumococci [21–23], some of which may be undetected at sampling time and not all of which may be affected by the vaccine. Given these complexities, design of an RCT for a new vaccine involves challenging questions of choosing the best population and inclusion criteria to improve the chances of seeing a real effect of the vaccine, choosing at what time after vaccination to measure carriage, and estimating power and sample size requirements.

Mathematical transmission modeling [15] and simulations [24–26] have been used to assist in the design of intervention trials for infectious diseases. These approaches have been needed, and useful, because standard assumptions about the magnitude of effect size and predictable event rates in controls are often not met in the setting of a transmissible pathogen, particularly when accounting for complexities like those mentioned above.

An inactivated whole cell pneumococcal (wSP) vaccine has recently been manufactured under Good Manufacturing Practices [27] and has been employed in dose-finding, immunogenicity, and safety studies in Kenyan adults and toddlers (clinicaltrials.gov NCT02097472) [28]. Although not powered for efficacy evaluation, this study was extended to evaluate nasopharyngeal carriage in toddlers participating in the trial. Based on murine data, it is believed that the primary impact of such a vaccine is to hasten the development of T-cell-mediated immunity to colonization, thereby reducing the duration of carriage episodes [17,29]. To aid in evaluating the results of this study and in planning future, larger studies, we undertook simulation modeling of such a trial in different age groups and settings to answer several questions:

1. What is the relationship between the amount of immune enhancement such a vaccine produces and the size of the effect on carriage prevalence in a setting similar to the Kenyan trial?
2. How does this relationship depend on the age of the trial participants (which affects their level of immunity at baseline, as well as their exposure to transmission during the trial), and on the intensity of transmission in the population (which affects the rate at which immunity develops in both vaccine recipients and controls)?
3. What are the implications for the sample size required to detect a particular effect size?
4. Which choice of setting, age group, and time from vaccination to carriage measurement will be most powerful in detecting various levels of vaccine impact on hastening immune development?

## Results

### Sampling time and participant age strongly influence sample size

Our simulation study was based on a published individual-based model of pneumococcal transmission that incorporates many of the complexities described above [30]. To this model, we added the ability to simulate vaccine trials, and implemented an algorithm to fit parameters to carriage prevalence data. The wSP vaccine was modeled as accelerating the exposure-dependent development of non-serotype-specific immunity against carriage duration, i.e. vaccination was immunologically equivalent to having cleared more colonizations. Three possible vaccine efficacies were considered: 3, 5, or 10 “colonization equivalents” (“c.e.”), which correspond, respectively, to an additional 26%, 39%, or 63% reduction in carriage duration. We assumed a minimum carriage duration of 20 days, and so reductions in duration affect the duration of carriage beyond the first 20 days. Trial participants in the model were vaccinated once, either as infants, at 60 days of age, or as toddlers, at 360 days, and the vaccine was assumed to be effective immediately upon receipt. Simulated trials took place in two settings that differed in their transmission intensity: the higher transmission setting had an under-five carriage prevalence of 66%; the lower transmission setting, 55%.

For the higher transmission setting, we ran 50 simulations of the vaccine trial using different random seeds and recorded the carriage prevalence every month (defined as 30 days), starting from birth to 24 months after vaccination (**Fig 1**). For both infants and toddlers, all vaccine efficacies led to reductions in prevalence throughout the follow-up period. Higher efficacies resulted in greater reductions in carriage. However, that marginal benefit attenuated with time as both controls and vaccinees acquired more natural immunity from carriage episodes. Similar patterns were observed in the toddler trials, but with smaller reductions in prevalence (**Fig 2A-C**).

**Fig 1.**
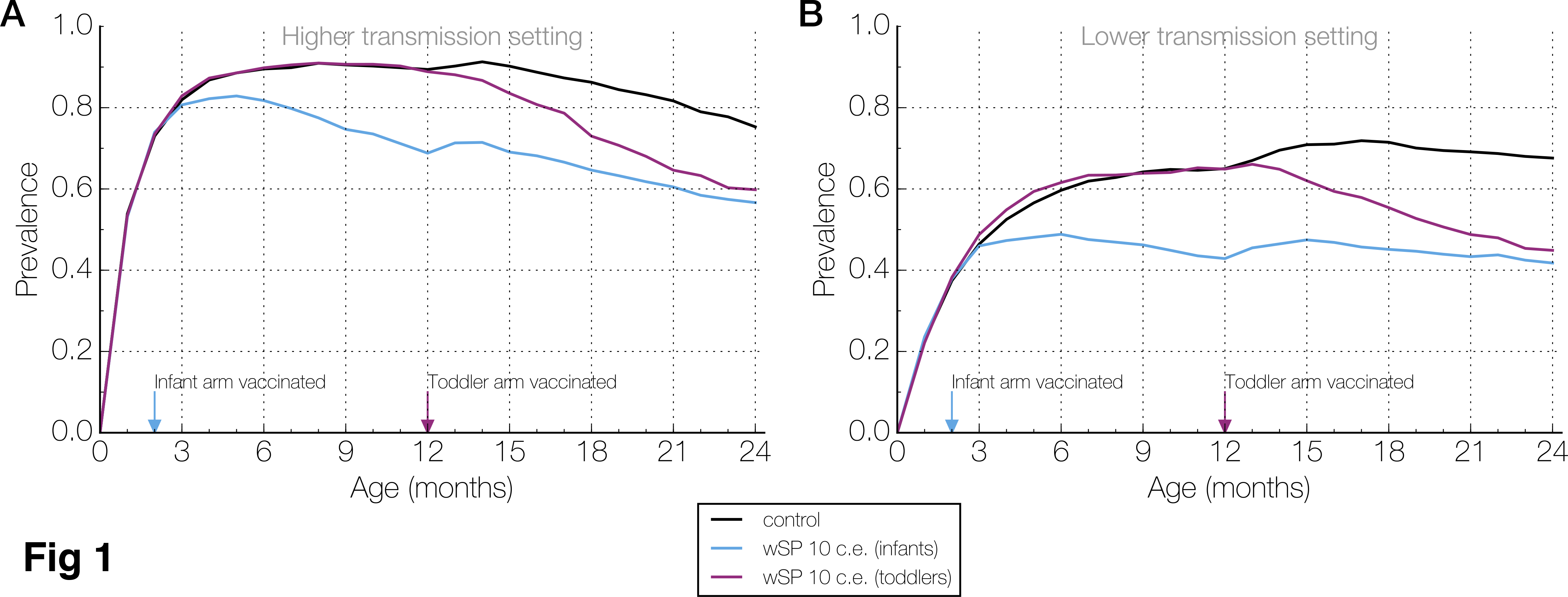
Age-specific carriage prevalences from representative simulation runs. (**A**) Carriage prevalences, sampled every month starting from birth, is shown for three arms – control (black), those vaccinated as infants (blue), and those vaccinated as toddlers (purple) – in a simulated trial in the higher transmission setting. Only the 10 colonization equivalent (c.e.) wSP vaccine efficacy is presented here. On the x-axis, two arrows indicate the age at which the vaccine was administered for the vaccinated arms. **(B)** Similar to (A), but for a simulated trial in the lower transmission setting.

**Fig 2.**
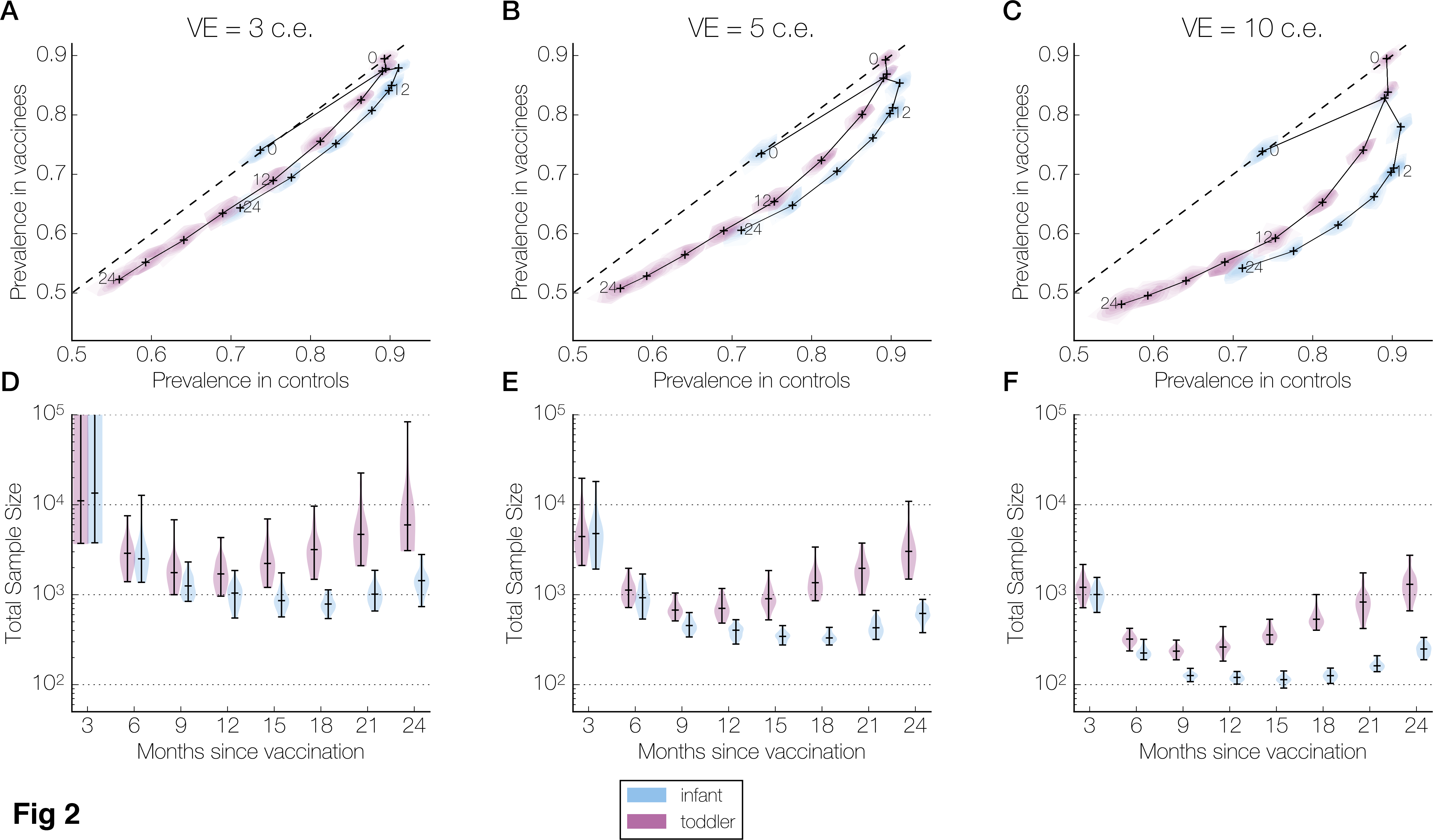
Prevalence and sample size over the follow-up period in the higher transmission setting. Panels are organized column-wise by vaccine efficacy: 3 colonization equivalents (c.e.), or 26% reduction in carriage duration (A, D); 5 c.e., or 39% (B, E); and 10 c.e., or 63% (C, F). Within each panel, results are presented separately for infants (blue) and toddlers (purple). **(A-C)** The joint kernel density estimate (see Methods) of the control and vaccine arm prevalences at each sampling time (every 3 months until 24 months post-vaccination) is shown as a contour map truncated by the convex hull of the simulated points, with the median values marked by a cross. These crosses are connected chronologically, and those corresponding to 0, 12, and 24 months post-vaccination are labeled. The dashed line indicates equal prevalences in the two arms. **(D-F)** The kernel density estimate of the total sample size (combined size of both samples) needed to detect a difference between control and vaccine arm prevalences at each sampling time (assuming 80% power, 5% type I error rate, balanced arms). The horizontal bars in each violin plot indicate the minimum, median, and maximum values across all simulations. In (D), the maximum sample sizes for infants and for toddlers at 3 months post-vaccination are greater than one million and not shown.

For the infants, the prevalence in the control and vaccine arms followed non-monotonic trajectories over the course of the follow-up period. In the infants, the median prevalence in the control arms started at 74% at 2 months of age, peaked at 91% at 8 months of age, and then declined (**Fig 2A-C**, **Fig 1A**). The timing of the peak is consistent with previously reported data from Kilifi, Kenya [31]. In the vaccinated infants, the median prevalence peaked at the same time, at 8 months of age for the 3 c.e. vaccine efficacy, or slightly earlier, at 5 months of age for the 5 c.e. and 10 c.e. wSP vaccine efficacies (**Fig 2A-C**, blue). For the toddlers, who are vaccinated later in life at 12 months of age, the age-specific prevalence in both the control and vaccine arms steadily declined across the 24-month follow-up period (**Fig 2A-C**, purple).

From the joint trajectory of the control and vaccine arm prevalence over the follow-up period, we determined how the sample size required for a two-sample test of equal proportion varied with sampling time. We assumed a 5% type I error probability, 80% power, and balanced arms, and use the term “sample size” to refer to the combined size of both arms. In infants, for all vaccine efficacies, the median sample size decreased dramatically—almost ten-fold or more—in the period 3 to 9 months post-vaccination, plateaued, and then started increasing around 18 months post-vaccination. In toddlers, the median sample size over time was also U-shaped, reaching a minimum at 9 months post-vaccination before increasing (**Fig 2D-F**, purple). At virtually all sampling times and for all vaccine efficacies, the median sample size was larger in the toddler trials than in the infant trials (**Fig 2D-F**).

### Lower transmission intensity lengthens window of favorable sampling times

To examine the impact of transmission intensity in the population on carriage prevalence in the trial, we also ran 50 simulations of the vaccine trial in the lower transmission setting. As in the higher transmission setting, all vaccine efficacies resulted in reductions in carriage prevalence at all sampling times. The prevalence peak previously observed in infants was delayed, due to the slower acquisition of non-serotype-specific immunity in a lower transmission setting (**Fig 1**). Thus, the prevalence trajectories in controls and vaccinees followed non-monotonic trajectories in both infants and toddlers (**Fig 3A-C**). In the infant arms, the kink in the prevalence trajectory between 9 and 12 months post-vaccination was due to the change in age-specific contact patterns as the participants aged into the next age group (**Fig 3A-C**, **Table S1**).

**Fig 3.**
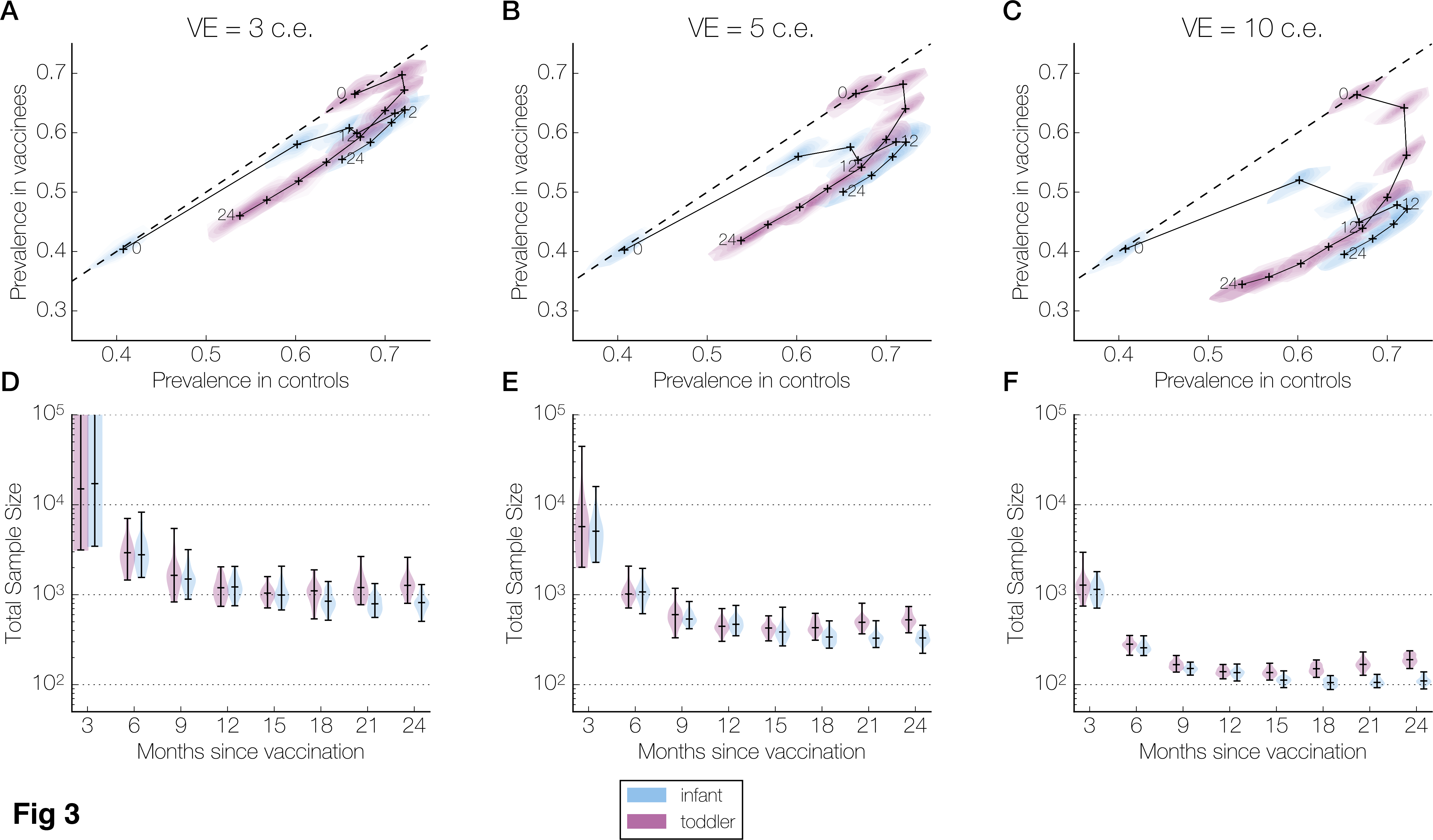
Prevalence and sample size over the follow-up period in the lower transmission setting. Panels are organized column-wise by wSP vaccine efficacy: 3 colonization equivalents (c.e.), or 26% reduction in carriage duration (A, D); 5 c.e., or 39% (B, E); and 10 c.e., or 63% (C, F). Within each panel, results are presented separately for infants (blue) and toddlers (purple). **(A-C)** The joint kernel density estimate (see Methods) of the control and vaccine arm prevalences at each sampling time (every 3 months until 24 months post-vaccination) is shown as a contour map truncated by the convex hull of the simulated points, with the median values marked by a cross. These crosses are connected chronologically, and those corresponding to 0, 12, and 24 months post-vaccination are labeled. The dashed line indicates equal prevalences in the two arms. **(D-F)** The kernel density estimate of the total sample size (combined size of both samples) needed to detect a difference between control and vaccine arm prevalences at each sampling time (assuming 80% power, 5% type I error rate, balanced arms). The horizontal bars in each violin plot indicate the minimum, median, and maximum values across all simulations. In (D), the maximum sample sizes for infants and for toddlers at 3 months post-vaccination are greater than one million and not shown.

As in the higher transmission setting, the total sample size decreased substantially in the period 3 to 9 months post-vaccination, and reached similar minimums. In the infant arms, the total sample size remained close to the minimum until the end of the 24-month follow-up period. In the toddler arms, the median sample size increased slightly near the end of the follow-up period. However, this rebound was considerably smaller than in the higher transmission setting, and the median sample size at 24 months post-vaccination was roughly five-to six-fold smaller. The sample sizes for the infant and toddler arms were more similar than in the higher transmission setting, particularly for later sampling times (**Fig 3D-F**).

## Discussion

Using a computational, individual-based transmission model of pneumococcal carriage, we estimated that a vaccine that enhances the immune response by an amount corresponding to 3, 5, or 10 carriage episodes could reduce age-specific carriage prevalence up to 7%, 10%, and 17%, respectively, compared to control in a setting similar to that of the wSP vaccine trial in Kenya, but that the magnitude of the reduction would depend strongly on the age at which participants were sampled. We found, however, that larger reductions could be observed if the same trial were performed in infants, in a lower-transmission setting, or both. Altogether, this analysis indicated that an infant trial conducted in a lower-transmission setting for a vaccine simulating 3, 5, or 10 exposures could be adequately powered with fewer than 800, 330, or 110 participants respectively, if the sampling window were chosen to be 15 to 24 months post-vaccination. Suboptimal choices of setting, age group, and sampling time could multiply the required sample size by a factor of ten or more.

The individual-based computational model [30] on which our work is based was originally used to explain serotype diversity and explore serotype replacement following the introduction of conjugate vaccines. With modifications, this model is also well suited to address our modeling questions, because it incorporates many processes, epidemiological and immunological, that complicate the relationship between the efficacy of a vaccine believed to reduce carriage duration but not risk of acquisition, and its effect on carriage prevalence. Our extensions—an algorithm to fit the model to specific epidemiological settings and the ability to randomize trial participants to different vaccine interventions—allow this model to be used for vaccine trial planning.

Our simulated vaccine trials show that sampling time and participant age greatly influence the number of participants needed to detect a protective effect of a vaccine whose effect is accelerating the development of immunity against carriage duration, as the wSP vaccine and perhaps other protein-based vaccines targeting carriage are expected to do. Across different combinations of vaccine efficacies and participant ages, the required sample size reached a minimum approximately 9 months post-vaccination before rebounding in later months. This favorable sampling time is consistent with simulation results by Scott et al., who explored similar questions, but more generally and for vaccines whose primary effect is on acquisition rather than duration, and using a compartmental transmission model [15]. This timing is also consistent with what Auranen et al., who explored pneumococcal trial design questions with a Markov transition model, suggest: waiting at least twice the average carriage duration after immune response before sampling [32].

In our simulations, the U-shaped trajectory of sample size over the follow-up period indicates a window of favorable sampling times, when the sample size is relatively small as compared to earlier or later. We found that sample sizes are lower, and the favorable window longer, when trial participants were younger, and when the transmission level was lower. In these scenarios, natural immunity is weaker initially or develops more slowly, and thus immune enhancement by the vaccine is more apparent. This intuition is what our simulation study attempts to quantitate, in terms of sample size, for different trial conditions.

Certain model assumptions may affect our conclusions. Our formulation of vaccine efficacy requires estimating the acquisition rate of exposure-dependent immunity. Direct estimates of vaccine efficacy against carriage, when they become available, can be used instead. We assume that the vaccine shortens only future carriage episodes, but not ones already present at the time of vaccination. Since the intrinsic duration of the fittest serotype is five months, this assumption would delay the vaccine’s effect on carriage prevalence, and thus, our reported favorable sampling times. This delay would affect infants more than toddlers, as they are more immunologically naïve and experience longer carriage durations. Auranen et al., in their study, report that sampling time is determined by the rate of clearance rather than rate of acquisition, which reinforces the importance of determining whether a vaccine accelerates the clearance of pre-existing carriage episodes [32]. Another important assumption is that exposure, rather than age alone, is responsible for the progressive shortening of carriage episodes as an individual gets older. If immune maturation due to calendar age, rather than or in addition to increased exposure, actually reduces carriage duration, then that would bolster the case for younger trial participants. Regardless of age at vaccination, the favorable sampling windows will likely be shortened as well. Our simulation framework can be easily updated should future evidence suggest revisiting these assumptions.

In its current form, our current simulation framework is already adaptable enough to examine a variety of scenarios. The ability to tailor simulations to specific settings is particularly useful—vaccine trials take place in countries with different age and serotype distributions, and Phase I/II and Phase III trials of the same vaccine may be conducted in the different locations. While we present results for a vaccine against carriage duration, we can also model vaccine protection against acquisition, and specify whether a vaccine effect is serotype-specific. The analysis presented here can be easily repeated, without changes to the source code, for trials involving polysaccharide conjugate vaccines, which protect against acquisition [4] and whose protection is serotype-specific [10], and novel vaccines with both polysaccharide and protein antigens [33], which may elicit a combination of serotype-specific and cross-reactive responses against carriage. The general population can also be vaccinated. Hence, our framework can be used to simulate trials—such as those comparing dosing schedules—that take place in countries with existing vaccination programs. In addition to planning future trials, our simulation framework can be used to examine completed trials. For completed trials with carriage endpoints that have not found a statistically significant vaccine effect, such as a recent phase II trial of a protein and polysaccharide-based vaccine in Gambian infants [33], simulation studies such as this can help assess whether inadequate power is a compelling explanation.

The analysis presented in this paper does not consider the effect of vaccination on carriage density or other factors (apart from duration) that would affect the infectiousness of a person who is vaccinated yet still becomes colonized. More generally, we do not consider the impact of vaccination on transmission at all in our simulations: simulated trial participants are computationally isolated from other hosts to approximate an individually randomized trial in which the participants are a negligible fraction of the population. However, our current framework can also simulate roll-outs of vaccination programs in the simulated population, where there is transmission between individuals, thus allowing the indirect effect of vaccination to be included. Vaccines with direct effects against transmissibility, possibly via reducing bacterial density in the nasopharynx, can be incorporated into our framework as well, with minimal modifications to the source code.

## Methods

### Mathematical model

#### Pneumococcal transmission dynamic model

This simulation study was based on a published individual-based model of pneumococcal carriage that incorporates many of the complexities relevant to our modeling questions [30]. Briefly, hosts are exposed to and can be colonized by multiple serotypes through age-specific contact with others. Serotypes differ in their mean duration of colonization in a naive host (“intrinsic duration”), which ranges from 20 to 150 days [19,20], and in their ability to prevent other strains from colonizing the same host. These phenotypes are positively correlated—i.e. fitter serotypes have longer intrinsic durations and are more likely to prevent concurrent colonizations—through their dependence on a serotype-specific fitness parameter. Hosts acquire immunity through colonizations. Clearing a colonization results in serotype-specific (anti-capsular) immunity that reduces risk of acquisition of the same serotype. Each clearance, of any serotype, enhances nonserotype-specific immunity that reduces the mean duration of carriage episodes.

#### wSP vaccine effect

The wSP vaccine was modeled as accelerating the acquisition of non-serotype-specific immunity that reduces carriage duration. As in Cobey et al. [30], the duration of a carriage episode is drawn from an exponential distribution with a mean given by

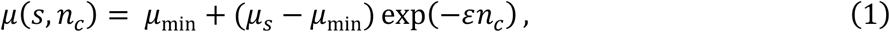

where *s* is the serotype carried, *n_c_* is the number of cleared carriage episodes (of any serotype), *µ*_min_ is the minimum mean duration, and *µ_s_* is the intrinsic duration of serotype *s*. The exposure-dependent development of non-serotype-specific immunity is captured in the exponential decay term in Equation 1. Each cleared colonization is immunizing, but with diminishing returns, and brings the mean duration closer to the minimum mean duration. For a vaccinated individual, the mean duration is given by

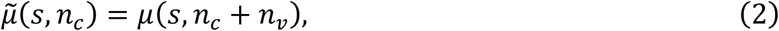

where *n_v_* is a positive constant characterizing the strength of the vaccine effect. Thus, the wSP vaccine can be thought of as boosting the non-serotype-specific immunity by an additional *n_v_* cleared colonizations, and we can express its efficacy in terms of “colonization equivalents” or “c.e.” We considered three different values of *n_v_*: 3, 5, and 10. The duration of each carriage episode was determined at the time of colonization, and hence, the vaccine did not affect colonizations already present on the day of vaccination. For simplicity, we assumed that full efficacy is achieved immediately upon receipt of a single dose.

#### Vaccine trials

To the original transmission model, we added the ability to simulate vaccine trials. Each trial arm was characterized by the number of participants, the enrollment date, and the vaccine and dose schedule used. In our implementation, trial participants were semi-isolated from the population: their demographics were tracked separately and their colonizations do not contribute to the force of colonization for the main population, but their exposures and risk of colonization were equivalent to those of the same age in the main population. This implementation design ensured that their colonization histories remain representative of participants within the main population, while affording two advantages: 1) We can have an arbitrarily large number of trial participants without skewing the epidemiological dynamics of the population, and 2) participants can be “enrolled” simply by birthing them into the simulation, without skewing the age structure of the population. Alternatively, we could have achieved these properties by simulating a large enough population such that the trial participants are a negligible fraction and thus do not create appreciable herd immunity in the population—the case in most real-world individually-randomized vaccine trials. However, that approach would have been considerably more computationally intensive.

#### Simulations

Simulations were initiated with hosts of different ages and no colonizations. The number of hosts was kept constant throughout a simulation. Every simulation was run first for 50 years to allow the age distribution of the population to stabilize, after which colonizations were seeded in the population and the simulation was run for another 50 years to allow the epidemiological dynamics to equilibrate. At this point, the simulated vaccine trial was initiated. For simplicity, all participants were birthed into the trial on the same calendar day. To reduce sampling noise, each trial arm had 5000 participants, 100-fold more than the trial arms in the Kenyan wSP study [28]. The participants were followed for five years and the carriage prevalence in each trial arm was recorded every 30 days. These carriage prevalences were then used as “true prevalences” to calculate the sample size needed to compare between arms, based on a two-sample test for equal proportions and assuming a 5% type I error rate, 80% power, and balanced arms [34]. We use “sample size” to refer to the combined size of both arms. All combinations of vaccine efficacies (3, 5, 10 c.e. and control) and ages at vaccination (60 and 360 days) were represented in each simulated trial (for a total of 8 arms), allowing us to control for transmission in the main population when comparing between arms. For computational speed, the main population was set at 25 thousand individuals. For each parameter set, we conducted 50 simulations runs - enough so that trends could be distinguished from stochastic variation between simulations, but not too many as to require an unreasonable amount of computation time. The model was implemented in C++11 with Boost C++ libraries. Analysis of simulation results was performed using Python 2.7 and browser-based Jupyter interactive notebooks [35]. Smoothed distributions were estimated using Gaussian kernel density estimation as implemented in the SciPy and Matplotlib Python libraries [36,37], and visualized as a violin plots (1-dimensional) or contour plots (2-dimensional).

### Parameter choices

We considered two settings that differ in their transmission intensity. The higher transmission setting was chosen to approximate Kenya, the site of a recent dose-finding and safety study [28]. The age distribution of simulated hosts was matched to that of Kenya’s population in 2015 [38], the second year of the study, which ran from April 2014 to December 2015. The age-specific mixing matrix was estimated from a social contact study in Kilifi, Kenya from 2011-2012 [39] and can be found in **Table S1**. The age structure in the model is described in more detail in **Text S1**. We fixed the non-serotype-specific immunity acquisition rate so the simulated age-specific carriage durations are consistent with the age-specific rates of clearance in Kenyan toddlers estimated by Abdullahi et al. [40] (**Fig S3**). The serotype fitness parameters were fit to serotype-specific carriage prevalences from a cross-sectional study in Kilifi from 2006 to 2008 [31], before the introduction of the conjugate vaccine PCV10. We chose to fit using only pre-PCV10 data. Trying to reproduce changes in serotype distribution due to PCV10 would have introduced additional complications, while being unlikely to yield further insight into our modeling questions given that the wSP vaccine is expected to act in a serotype-agnostic manner [41]. A mathematical description of the fitting algorithm can be found in **Text S2** and the fitted serotype fitness parameters are listed in **Table S2**.

For the lower transmission setting, we used a smaller overall contact rate, so the simulated carriage prevalence at 12 months of age resembles preliminary estimates from a study in Indonesia [42], the proposed site for a follow-up wSP vaccine efficacy trial (**Fig S3**). To facilitate comparisons between settings, we kept the same age distribution, age-specific mixing pattern, and fitness parameters used in the higher transmission setting. A summary of the model parameters and their values can be found in **Table 1**.

**Table 1.** Selected^1^ model parameters.

| Symbol<sup>2</sup> | Description | Value(s) | Refs |
| --- | --- | --- | --- |
| **Demographic** |  |  |  |
| $N(0)$ | Number of hosts (in thousands) | 25 | Main text |
| - | Maximum age (years) | 101 | [38], **Text S1** |
| - | Lifespan distribution | **Fig S1** | [38], **Text S1** |
| $\alpha$ | Age-specific mixing weights | **Table S1** | [39], **Text S1** |
| **Epidemiological** |  |  |  |
| - | Minimum intrinsic duration (days) | 20 | [20] |
| - | Maximum intrinsic duration (days) | 150 | [19] |
| $\kappa$ | Minimum carriage duration in any host (days) | 20 | [20] |
| $\mu_{\max}$ | Maximum competitive exclusion | 0.25 | [20] |
| - | Serotype fitness parameters | **Table S2** | **Text S2** |
| $\epsilon$ | Acquisition rate of non-serotype-specific immunity | 0.1<sup>#</sup> | [40] |
| $\beta$ | Overall contact rate (contacts per day per host) | 0.1 or 0.13<sup>&</sup> | **Text S2** |
| **Vaccine trial** |  |  |  |
| $a_v$ | Age at vaccination (days) | 60 or 360 | Main text |
| - | Vaccine efficacy (colonization equivalents) | 26%, 39%, or 63%<sup>‡</sup> | Main text |
| - | Number of participants per arm (in thousands) | 5 | Main text |

<sup>1</sup> Parameters adequately described in Cobey and Lipsitch's paper [30] are not repeated here. Parameters in this table either have new values, or are newly introduced.
<sup>2</sup> The symbol used in Cobey and Lipsitch's paper [30], or "-" if no symbol was used or if the parameter is new.
<sup>#</sup> Chosen such that age-specific carriage duration is consistent with previous clearance rate estimates.
<sup>&</sup> Fit to carriage prevalence in Indonesia and Kenya, respectively, and the only parameterization difference between the lower and higher transmission settings used in this paper.
<sup>‡</sup> Reduction in carriage duration, in addition to that due to natural immunity. Corresponding to the amount of immune enhancement from 3, 5, or 10 additional carriage episodes.

### Sensitivity analyses

To isolate the effect of transmission intensity in our main analyses, we had used the same age-specific mixing pattern-based on Kenya contact survey data [39]–in both the higher and lower transmission settings. Real-world vaccine trials, however, will take place in the context of different mixing patterns, or may be planned in the absence of reliable social contact data. To examine the robustness of our findings to the pattern of age-specific mixing, we repeated our analyses assuming random mixing between individuals, i.e., equal contact rate for all pairs of individuals. We re-fit the model to the observed Kenya carriage survey data [31], and ran a set of 50 simulations. With a random mixing pattern, there was a slightly higher carriage prevalence in trial participants during the first two years of follow-up. However, the total sample sizes, in both magnitude and trend across sampling time, remained similar to those from the main analyses (**Fig S4**, **Fig 2**).

Other potential sources of bias were the population and trial arm sizes. In the main analyses, we chose values that were small enough to allow simulations to finish reasonably quickly, and reduced the effect of simulation variability by running multiple simulations and considering sample median. To assess whether the sample median may be biased, we performed univariate sensitivity analyses of the population and trial arm size. Specifically, within the higher transmission setting, we varied population size between 10K, 25K, and 50K individuals (not including trial participants), with the trial arm size fixed at 5K. We also varied the trial arm size between 2.5K, 5K, or 10K participants, with the population size fixed at 25K. Note that the middle values, a population size of 25K and a trial arm size of 5K, were the ones used in the main analyses. Twenty-five simulations were run for each set of parameter values. Varying the population and varying the trial arm size did not appreciably alter the sample median of the simulated carriage prevalences (**Fig S5**). Larger population sizes led to smaller variability between simulations, which is expected given the stochastic nature of transmission in the model (**Fig S5A, B**). Larger trial arm sizes did not reduce variability, suggesting that the epidemiological dynamics in the general population are driving the variability in the trial arm prevalences, at least for the trial arm sizes examined (**Fig S5C, D**).

### Code repository

C++11 code for fitting and simulating the individual-based model can be found in the Github repository linked here: [will include link before publication]

## S1 Text Model age structure

The age distribution and age-specific contact rate of hosts is important to consider in pneumococcal transmission modeling, since carriage prevalence varies with age [1,2], as does frequency of contact with other age groups [3,4].

The age distribution of the simulated hosts was matched to the 2015 age distribution in Kenya, based on data from the United Nations World Population Prospects [5]. The number of simulated hosts was constant, and for a fixed-sized population, we can set its age distribution by choosing the correct lifespan distribution: For a simulated host, the probability of living exactly *n* years is calculated as the difference in the number of *n*-year old people and *n +* 1-year old people, divided by the total number of people. For this method to be valid, the age distribution must be monotonically decreasing, i.e. there cannot be more people in an older age class as compared to any younger age class. This is the case for Kenya’s age distribution in 2015. The World Population Prospects data was given in 5-year age classes, which we linearly interpolated to obtain 1-year age classes. The oldest age class in the data was 100 years or greater; in our model, we assume that the maximum lifespan is 101 years.

We derived age-specific mixing weights from social contact data collected in Kilifi, Kenya from 2011 to 2012 by Kiti et al [3]. Specifically, normalized the age group-specific average number of contacts per day by the size of the contacting age group and the size of the contacted age group. Since we fit the overall contact rate, for simplicity, we scaled the mixing weights so the maximum is 1. The weights used can be found in **Table S2**.

## S2 Text Model fitting algorithm

To simulate specific epidemiological settings, we implemented an algorithm that fit model parameters to given serotype-specific carriage prevalences, e.g. prevalences from survey data. In our model, the prevalence of each serotype is determined primarily by its fitness parameter and the overall contact rate shared by all serotypes. The fitness parameter can take values, possibly non-integral, from 1 to *n_s_*, the number of serotypes, Lower values correspond to better fitness. Lowering the fitness parameter results in two phenotypic changes—longer colonization duration and enhanced competitive ability— that both increase prevalence. Hence, there is a monotonic relationship between a serotype’s fitness parameter and its expected carriage prevalence, and this allows us to tune the fitness parameters in a straightforward manner.

The algorithm iteratively updates its estimate of the serotype fitness parameters. Let the current estimate at the start of iteration *k* be denoted by the vector 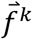, indexed by serotype. We run a simulation using 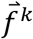. For serotype *s*, let 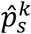 be its average prevalence over the last 25 simulation years, *p_s_* be its observed prevalence, and 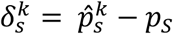 be the serotype-specific prevalence error. Based on this error, we update our estimate of the serotype’s fitness parameter according to:

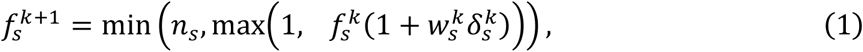

where the prevalence error is weighted by a factor 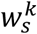 (**Fig S2A**). This factor is also updated iteratively, by comparing the prevalence error between the current and previous iteration. If the magnitude of the prevalence error is not decreasing enough between iterations, we increase the influence of the prevalence error in our updating of the fitness parameter, i.e. if 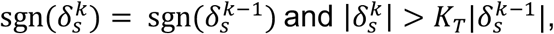 then 
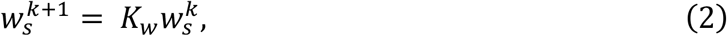

where *k_t_* is a positive constant and *k_w_* is a constant greater than 1. On the other hand, if the magnitude of the prevalence error decreased enough between iterations, or if it has changed signs and has become larger in magnitude, then we reduce the influence of the prevalence error in our update, i.e. if 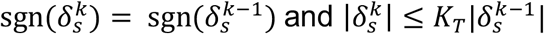 or 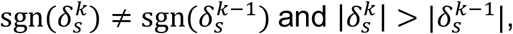 then 
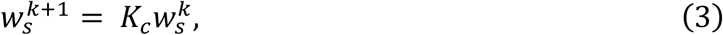
where *k_c_* is positive constant less than 1. By adjusting 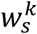 between iterations, we facilitate convergence of the fitness parameters: Equation (2) allows the algorithm to make larger adjustments when it is progressing too slowly, and Equation (3) causes the algorithm to be more cautious it is progressing quickly, or when the simulated prevalences start to oscillate around the observed prevalence. The latter is an indication that we are close to the optimal value for the fitness parameter—since the simulations are stochastic, we would not expect a properly fitted model to reproduce the observed prevalence exactly, but rather a distribution of simulated prevalences centered on the observed prevalence (**Fig S2B**).

This algorithm attempts to fit all serotype-specific prevalences simultaneously. It assumes that adjusting the fitness parameter of one serotype does not affect the prevalence of another serotype. Since there is competition between serotypes for hosts, that assumption is not strictly true. Nevertheless, we find that in practice, the fitting algorithm is able to converge reasonably quickly, within 125 iterations when using a population size of 20,000.

There are *n_s_* observed serotype-specific prevalences we are fitting to, but *n_s_ + 1* parameters: the *n_s_* serotype fitness parameters and the overall contact rate. So that the model is not underspecified, we fix the fitness parameter for the fittest serotype to be 1, which corresponds to an intrinsic colonization duration of 150 days and a relative reduction of 0.25 in the risk of colonization by other strains. With one of the fitness parameter fixed, we are free to fit the contact rate. Let *β^k^* be the current estimate of the contact rate in iteration *k*. Let 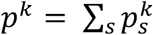 be simulated carriage prevalence during iteration *k*, *p* = Σ*_s_p_s_* be the observed carriage prevalence, and *δ*^*k*^ = *p^k^* – *p* be the total prevalence error. The update equation for *β* is similar to that of the fitness parameters:

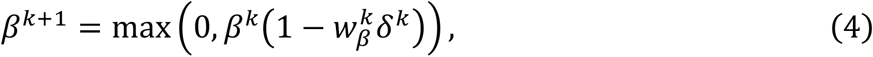

where *w_β_* is a positive constant. As before 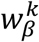 is updated as well, in the same fashion as described above for 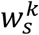, but with updating rules based on the *δ*^*k*^ rather than 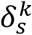.Parameters related to the fitting algorithm are summarized in **Table S1**.

## Supporting Information

**Text S1. Model age structure.** Derivation of the lifespan distribution and age-specific contact weights used in the model.

**Text S2. Model fitting algorithm.** Mathematical description of the algorithm used to fit the transmission model to carriage prevalence data.

**Table S1.**
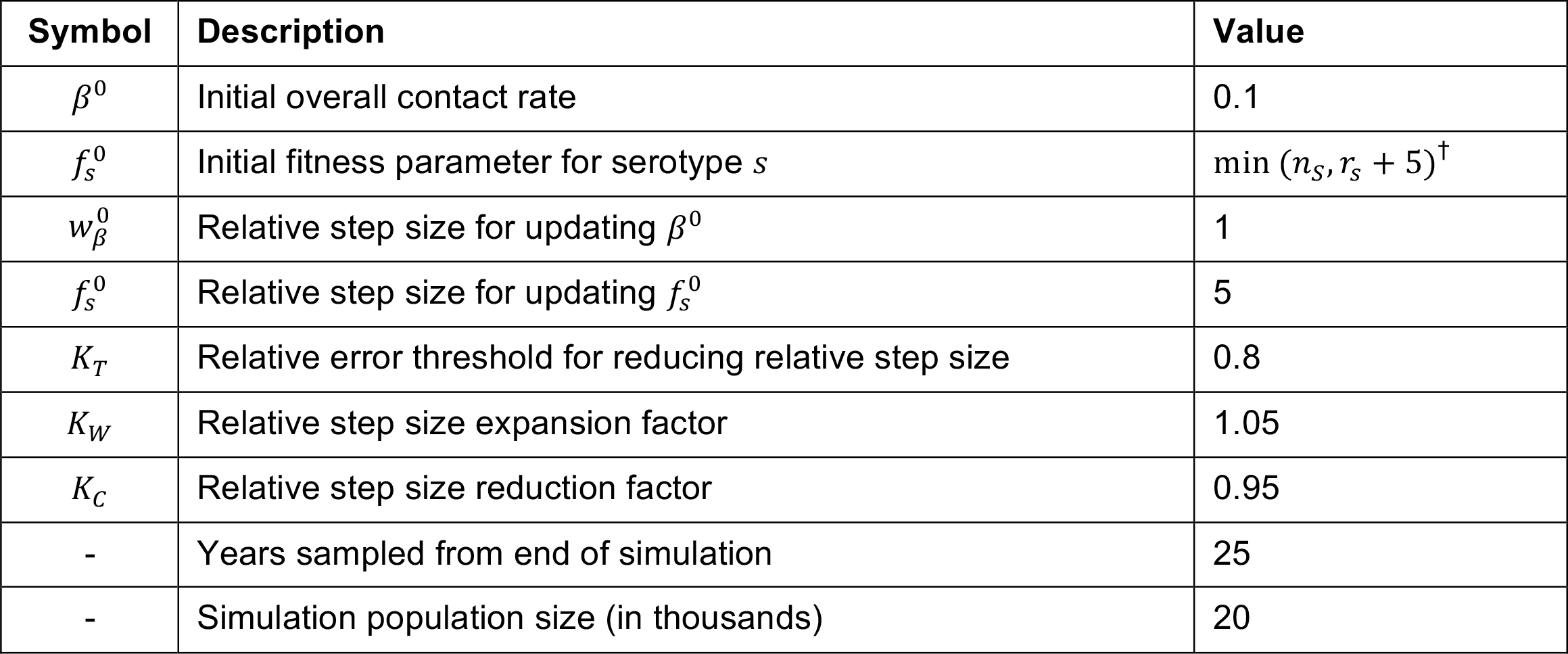
Age-specific mixing matrix.

| Symbol | Description | Value |
| --- | --- | --- |
| $\beta^0$ | Initial overall contact rate | 0.1 |
| $f_s^0$ | Initial fitness parameter for serotype $s$ | $\min(n_s, r_s + 5)^{\dagger}$ |
| $w_{\beta}^0$ | Relative step size for updating $\beta^0$ | 1 |
| $f_s^0$ | Relative step size for updating $f_s^0$ | 5 |
| $K_T$ | Relative error threshold for reducing relative step size | 0.8 |
| $K_W$ | Relative step size expansion factor | 1.05 |
| $K_C$ | Relative step size reduction factor | 0.95 |
| - | Years sampled from end of simulation | 25 |
| - | Simulation population size (in thousands) | 20 |

**Table S2.** Fitted serotype fitness parameters.

| **Age**<br>**(years)** | **<1** | **1-5** | **6-14** | **15-20** | **21-50** | **>50** |
| --- | --- | --- | --- | --- | --- | --- |
| **<1** | 0.1391 | 0.3739 | 0.4017 | 0.2938 | 0.4015 | 0.2566 |
| **1-5** | 0.3739 | 0.6283 | 0.5460 | 0.2844 | 0.3561 | 0.2517 |
| **6-14** | 0.4017 | 0.5460 | 0.8344 | 0.4775 | 0.3067 | 0.2304 |
| **15-20** | 0.2938 | 0.2844 | 0.4775 | 1.0000 | 0.4243 | 0.2877 |
| **21-50** | 0.4015 | 0.3561 | 0.3067 | 0.4243 | 0.7304 | 0.5665 |
| **>50** | 0.2566 | 0.2517 | 0.2304 | 0.2877 | 0.5665 | 0.5582 |

**Table S3.**
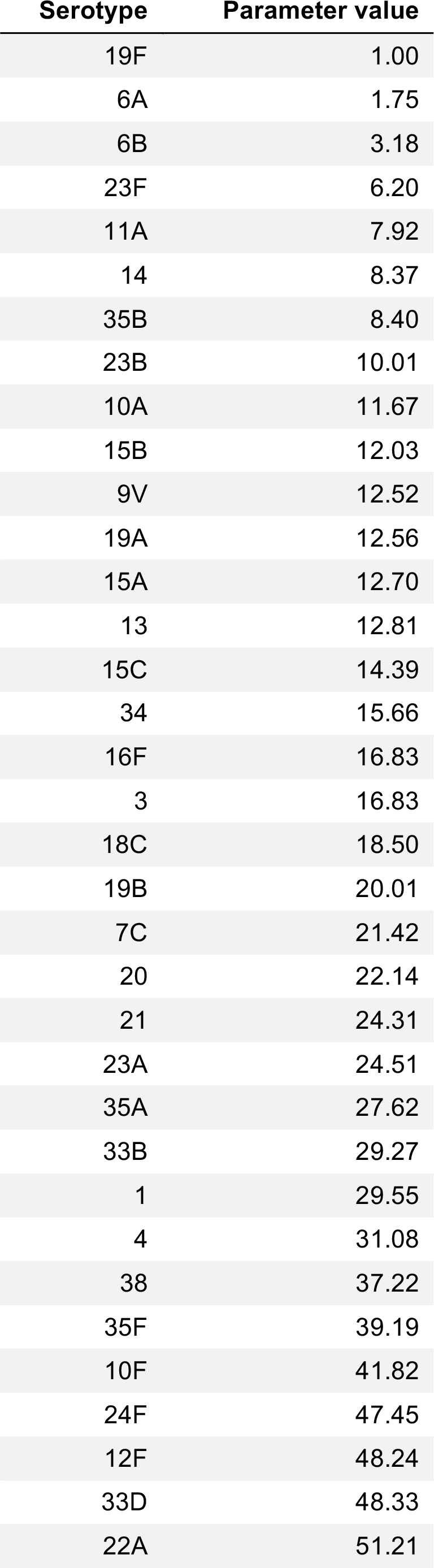
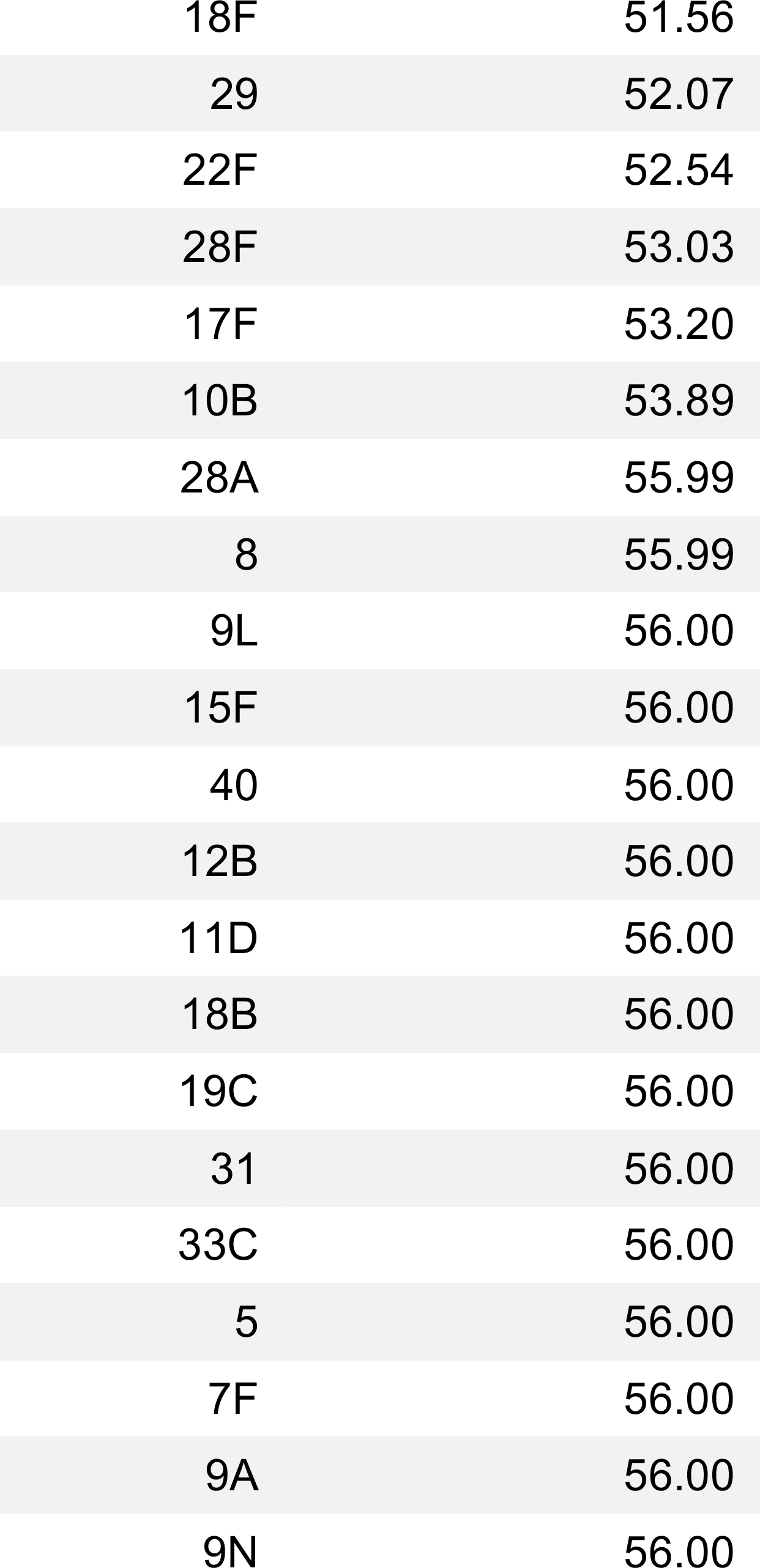
Parameters of the fitting algorithm.

| **Serotype** | **Parameter value** |
| --- | --- |
| 19F | 1.00 |
| 6A | 1.75 |
| 6B | 3.18 |
| 23F | 6.20 |
| 11A | 7.92 |
| 14 | 8.37 |
| 35B | 8.40 |
| 23B | 10.01 |
| 10A | 11.67 |
| 15B | 12.03 |
| 9V | 12.52 |
| 19A | 12.56 |
| 15A | 12.70 |
| 13 | 12.81 |
| 15C | 14.39 |
| 34 | 15.66 |
| 16F | 16.83 |
| 3 | 16.83 |
| 18C | 18.50 |
| 19B | 20.01 |
| 7C | 21.42 |
| 20 | 22.14 |
| 21 | 24.31 |
| 23A | 24.51 |
| 35A | 27.62 |
| 33B | 29.27 |
| 1 | 29.55 |
| 4 | 31.08 |
| 38 | 37.22 |
| 35F | 39.19 |
| 10F | 41.82 |
| 24F | 47.45 |
| 12F | 48.24 |
| 33D | 48.33 |
| 22A | 51.21 |
| 18F | 51.56 |
| 29 | 52.07 |
| 22F | 52.54 |
| 28F | 53.03 |
| 17F | 53.20 |
| 10B | 53.89 |
| 28A | 55.99 |
| 8 | 55.99 |
| 9L | 56.00 |
| 15F | 56.00 |
| 40 | 56.00 |
| 12B | 56.00 |
| 11D | 56.00 |
| 18B | 56.00 |
| 19C | 56.00 |
| 31 | 56.00 |
| 33C | 56.00 |
| 5 | 56.00 |
| 7F | 56.00 |
| 9A | 56.00 |
| 9N | 56.00 |

**Fig S1.**
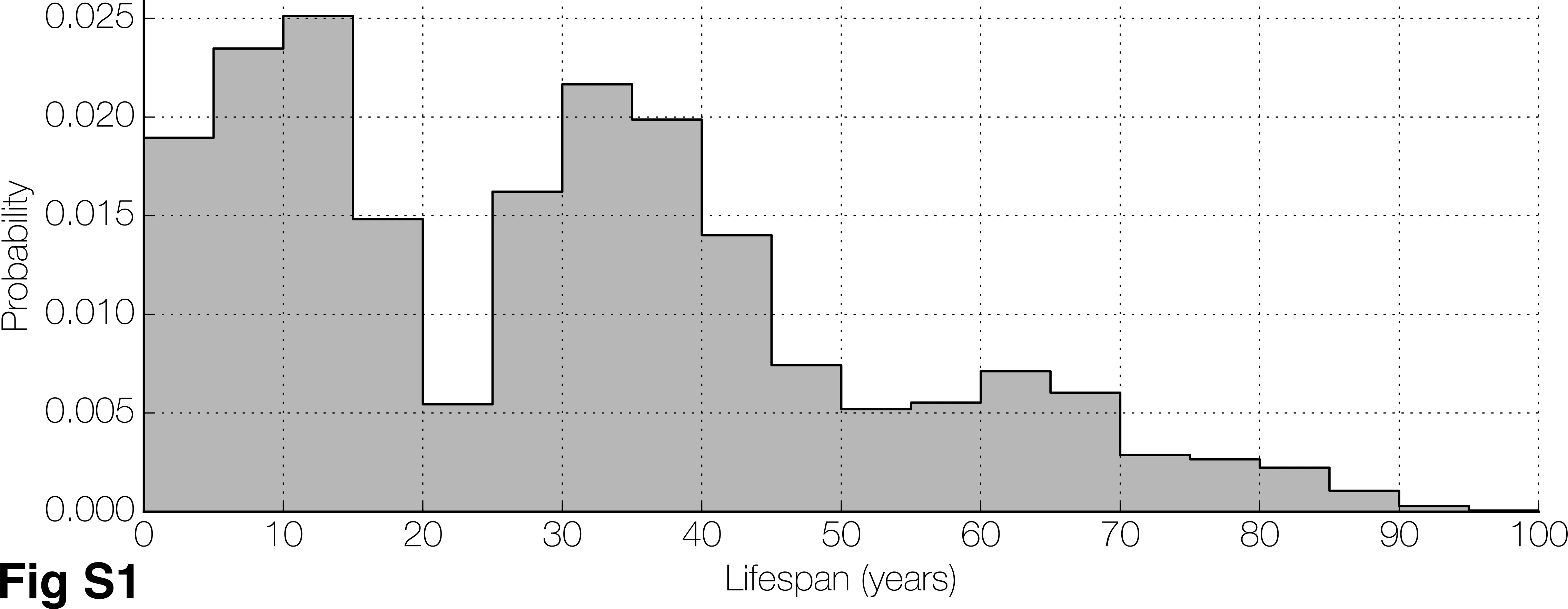
Lifespan distribution. The lifespan distribution used in all simulations. It is derived by assuming that the 2015 Kenya age distribution [38] is stable, i.e. no population growth. The step-wise nature of the distribution reflects the five-year intervals in the age distribution data.

**Fig S2.**
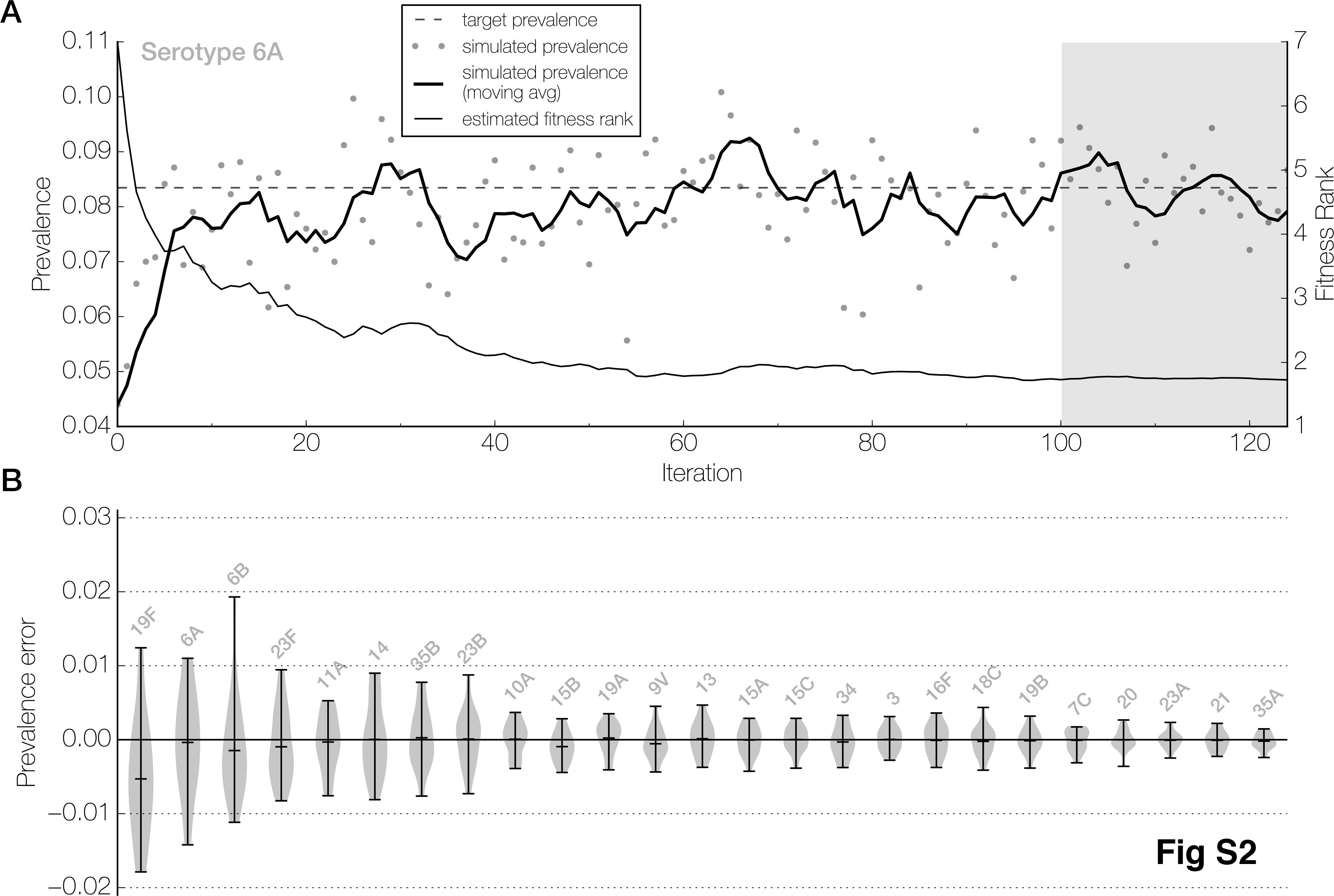
Estimation of serotype fitness parameters. **(A)** The fitting process for one representative serotype,6A. The evolving estimate of 6A’s fitness parameter (thin line, right y-axis) and 6A’s simulated prevalence (gray dots, left y-axis) is shown over the course of 125 iterations. Lower values of the fitness parameter correspond to a fitter phenotype. The moving average (thick line, n=5) of the simulated prevalences more clearly shows the trend of the simulated prevalences towards the target prevalence (horizontal dashed line). The light gray shaded region highlights the last 25 iterations, whose results are considered in (B). **(B)** One method of assessing the quality of the model fit. The distribution of prevalence errors (simulated minus target prevalence) in the last 25 iterations of the fitting process is shown for the top 25 serotypes (out of 56 total) by target prevalence (ranging from 9.96% for 19F to 0.53% for 35A). Each distribution is represented by a violin plot labeled by serotype name, and with horizontal bars marking the minimum, mean, and maximum values.

**Fig S3.**
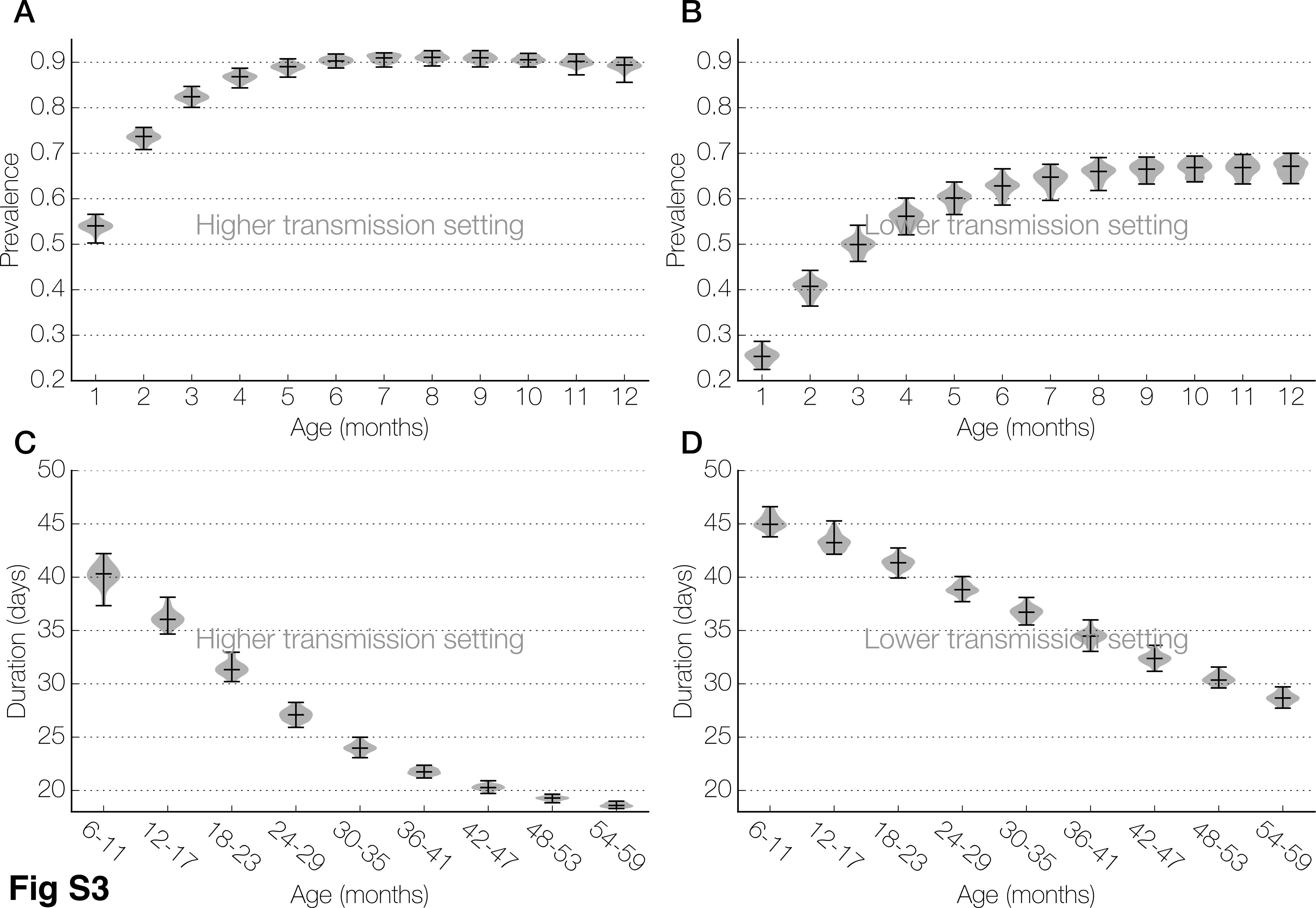
Age-specific carriage prevalence and duration. **(A, B)** Distribution of carriage prevalence in infants, by 1-month age categories, for the higher (A) and lower (B) transmission settings. (**C**, **D**) Distribution of carriage duration in infants and toddlers, by 6-month age categories, for the higher (C) and lower (D) transmission settings. Distributions are shown as violin plots, with horizontal bars indicating the minimum, median, and maximum values.

**Fig S4.**
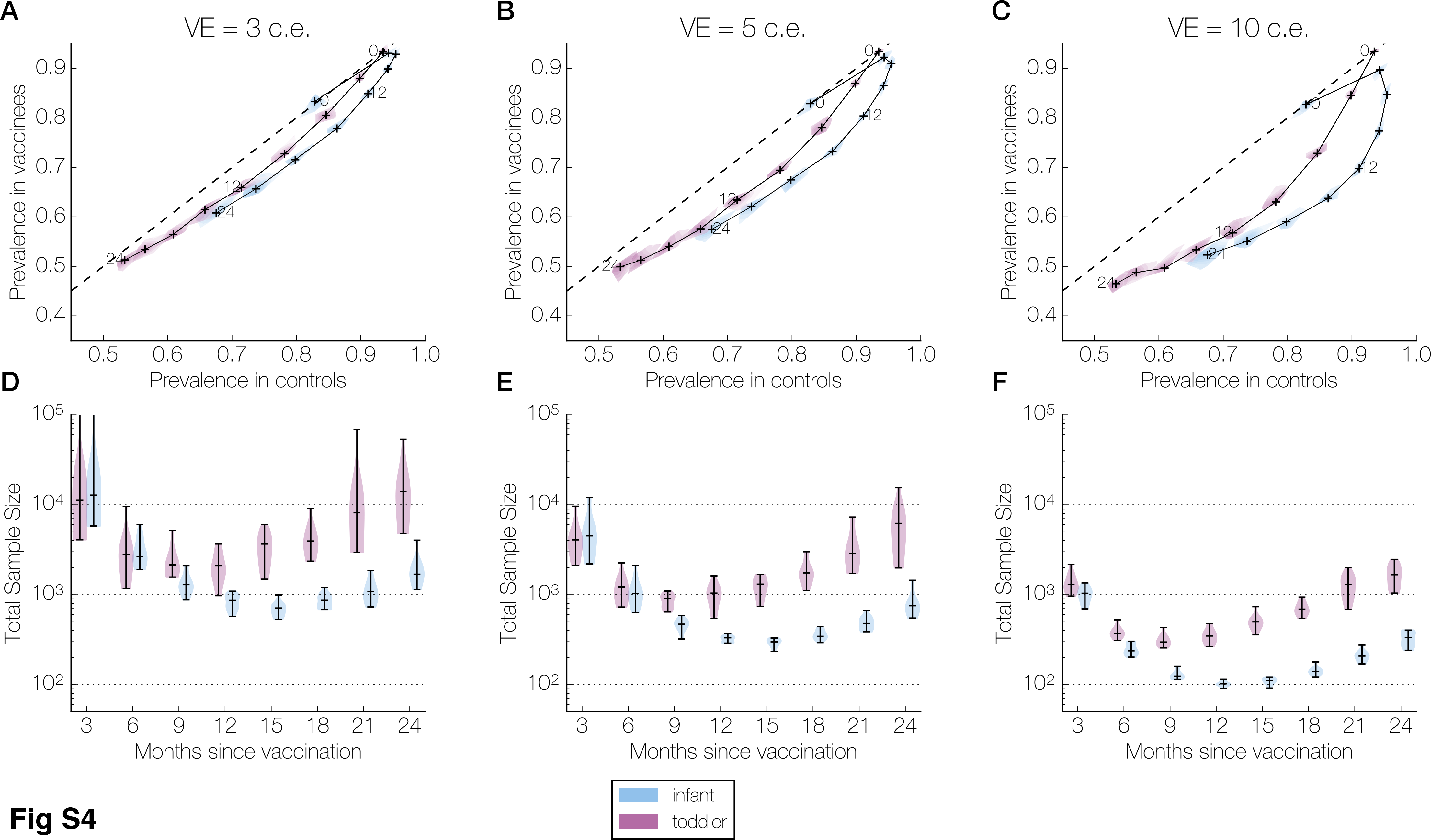
Prevalence and sample size over the follow-up period in the higher transmission setting, without age-structured mixing. Panels are organized column-wise by wSP vaccine efficacy: 3 colonization equivalents (c.e.), or 53% reduction in carriage duration (A, D); 5 c.e., or 71% (B, E); and 10 c.e., 92% (C, F). Within each panel, results are presented separately for infants (blue) and toddlers (purple). **(A-C)** The joint kernel density estimate (see Methods) of the control and vaccine arm prevalences at each sampling time (every 3 months until 24 months post-vaccination) is shown as a contour map truncated by the convex hull of the simulated points, with the median values marked by a cross. These crosses are connected chronologically, and those corresponding to 0, 12, and 24 months post-vaccination are labeled. The dashed line indicates equal prevalences in the two arms. **(D-F)** The kernel density estimate of the total sample size (combined size of both samples) needed to detect a difference between control and vaccine arm prevalences at each sampling time (assuming 80% power, 5% type I error rate, balanced arms). The horizontal bars in each violin plot indicate the minimum, median, and maximum values across all simulations. In (D), the maximum sample sizes for infants and for toddlers at 3 months post-vaccination are greater than one hundred thousand and not shown.

**Fig S5.**
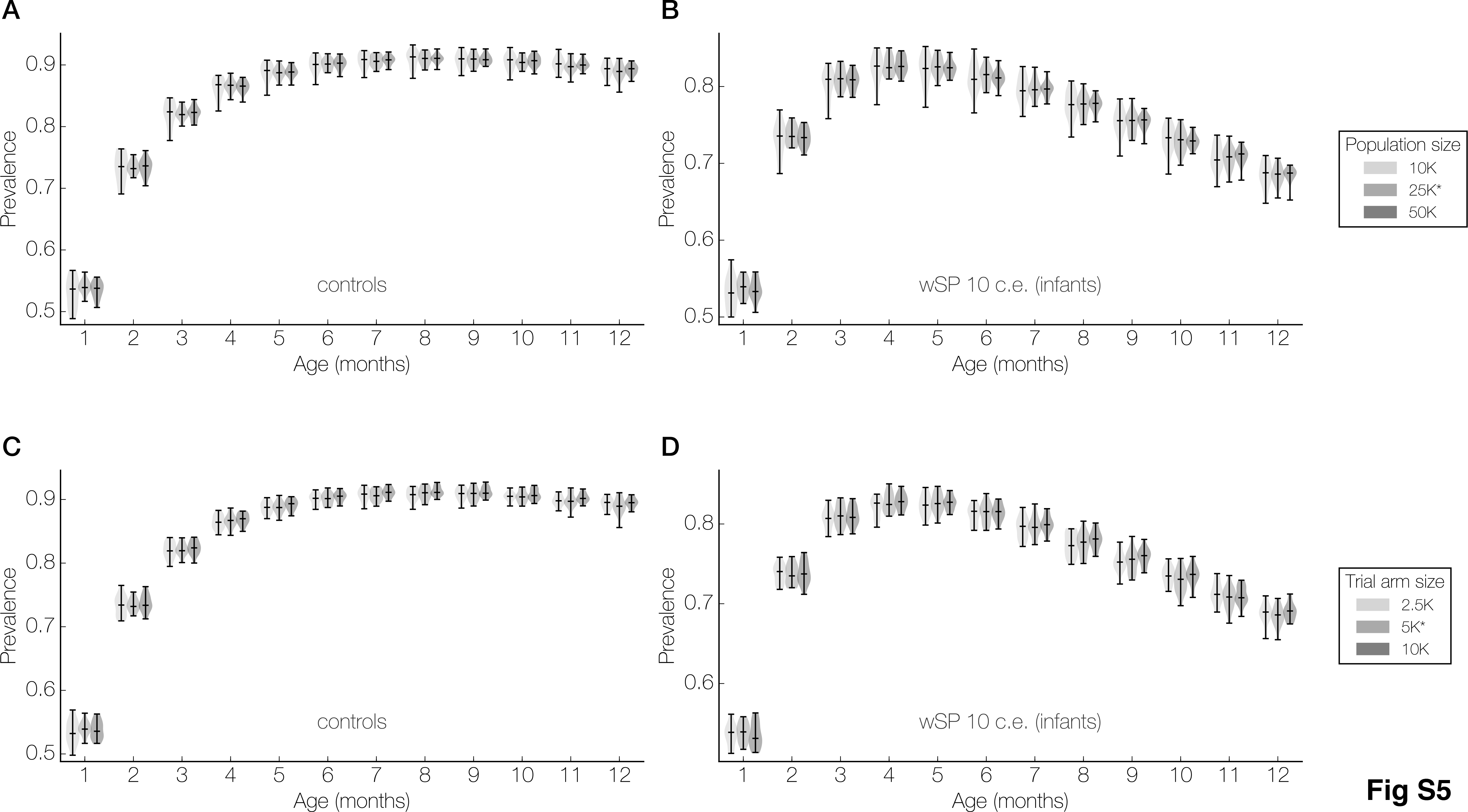
Population and trial arm size sensitivity analyses. **(A)** The age-specific prevalence in the control and wSP 10 c.e. (conferring an additional 92% reduction in carriage duration) infant arms for three different population sizes – 10K, 25K, and 50K individuals – with the trial arm size fixed at 5K participants. (**B**) The age-specific prevalence in the control and wSP 10 c.e. infant arms for three different trial arm sizes – 2.5K, 5K, and 10K participants – with the population size fixed at 25K. Each violin plot shows the distribution of prevalences across 25 simulations, with horizontal bars marking the minimum, median, and maximum values, and darker shades indicating larger population or trial arm sizes. The values used in the main analyses – a population size of 10K and a trial arm size of 5K – are marked with asterisks in the legends.

